# Males of a suboscine bird (*Ceratopipra mentalis*) exhibit individual distinctiveness in their contact call

**DOI:** 10.64898/2026.10.14.758842

**Authors:** Josephine T. Tagestad, Corey E. Tarwater, J. Patrick Kelley

**Affiliations:** Department of Zoology and Physiology, University of Wyoming, 1000 E. University Ave, Laramie, WY 82071, USA

**Keywords:** behavior, discriminant function analysis, lek, manakin, Neotropics, vocalizations

## Abstract

The ability to discriminate among individual conspecifics may play a role in competition, mate selection, and other social interactions. Studies of individual distinctiveness in birds have traditionally focused on oscines, owing to the assumption that the less vocally complex suboscines would be incapable of producing individually distinct vocalizations. Nevertheless, while studies remain limited, a growing body of literature indicates that suboscines are capable of producing individually distinct vocalizations. *Ceratopipra mentalis*, the red-capped manakin, is a lekking suboscine and an important seed disperser. This study investigates whether individual distinctiveness in the whistle-peep vocalization, a contact call between males, of *C. mentalis* exists by recording males at multiple leks in Panama. Using flexible discriminant function analyses, we found individual distinctiveness in temporal and frequency traits of the whistle-peep call structure. We were able to classify individuals with 81.1% accuracy, comparable to other studies of suboscines. These results provide further evidence that the absence of song learning and less complex syrinxes in suboscines do not preclude individual distinctiveness in vocalizations. While studies focus on territorial species, here we found individual distinctiveness in a non-territorial lekking species. We argue for the need to examine the role of vocalizations in mate selection and male-male competition in these species with intense sexual selection. Lastly, our results suggest that acoustic monitoring may be a useful path forward to obtain a rapid assessment of male population abundance in *C. mentalis* and potentially other suboscine species, critical given predicted changes in species with climate change and other human-induced impacts.

## INTRODUCTION

The ability of individuals to recognize members of their own species is important for social interactions, including territory defense and mate selection (Lambrechts and Dhondt 1995). Studies across taxa have demonstrated that many species are able to discriminate between conspecifics using individually distinct information contained in reliable sensory indicators of identity, such as sound or scent (Christensen and Radford 2018). For example, acoustic-based individual discrimination has been demonstrated in vervet monkeys (*Cercopithecus aethiops)*, Mexican free-tailed bats (*Tadaria brasiliensis mexicana)*, red deer (*Cervus elaphus)*, and Black-capped Chickadees (*Poecile atricapillus*) (Cheney and Seyfarth 1980, Balcombe 1990, Charlton et al. 2007, Hahn et al. 2017). In species that use vocal signals, individuals can typically be distinguished based on acoustic features in their vocalizations (Bard et al. 2002, Fitzsimmons et al. 2008, Clink et al. 2017, Gonázles-García et al. 2017).

Much research on vocal signals and individual acoustic distinctiveness has focused on birds (Naguib and Riebel 2006). However, until recently, studies of bird vocalizations and their functions focused almost exclusively on oscine (suborder Passeri) birds rather than suboscines (suborder Tyranni) (Irestedt et al. 2002, Moseley and Wiley 2013, Bruni and Foote 2014,).

Suboscine birds represent approximately 20% of the order Passeriformes, the largest avian order with more than 6,000 extant species worldwide (Sibley and Monroe 1990, Oliveros et al. 2019). Oscines are frequently assumed to be more vocally complex than suboscines, owing to the difference in syringeal complexity between the two groups (Ames 1971, Payevsky 2014, García et al. 2017). Oscine and suboscine birds also differ in vocal development, with oscines exhibiting song learning, while suboscine vocalizations are generally found to be innate (Kroodsma 1984, Baptista and Trail 1992, Touchton et al. 2014; see Kroodsma et al. 2013 for an exception).

Given these fundamental differences, individual vocal distinctiveness and the ability to recognize conspecifics (e.g., discrimination) based on vocalizations was traditionally thought to be exclusive to oscine species (Kroodsma 1984). However, recent studies have begun to overturn this assumption, as individual vocal distinctiveness has been demonstrated in vocalizations of Spotted Antbirds (*Hylophylax naevioides*), Alder Flycatchers (*Empidonax alnorum*), Acadian Flycatchers (*Empidonax virescens*), Screaming Pihas (*Lipaugus vociferans*), Peruvian and Yellow-breasted Warbling Antbirds (*Hypocnemis peruviana* and *Hypocnemis subflava*), Eastern Wood-Pewees (*Contopus virens*), Mexican Ant-thrushes (*Formicarius moniliger*), Eastern Phoebes (*Sayornis phoebe*), and Buff-throated Woodcreepers (*Xiphorhynchus guttatus*) (Bard et al. 2002, Lovell and Lein 2004, Wiley 2005, Fitzsimmons et al. 2008, Seddon and Tobias 2010, Clark and Leung 2011, Kirschel et al. 2012, Foote et al. 2013, Moseley and Wiley 2013).

Though these studies challenge the general assumption that only oscine birds exhibit individual distinctiveness, more investigation is needed across different suboscine species to determine if this ability is commonplace.

Manakins (family Pipridae) are suboscine frugivores that serve as essential seed dispersers in Neotropical ecosystems. Though vocal matching and microgeographic variation has been investigated in some manakin species (Trainer and McDonald 1995, Trainer and Parsons 2001), individual distinctiveness of vocalizations has yet to be examined in this diverse clade. In this study, we test whether the contact calls of red-capped manakin (*Ceratopipra mentalis*) males are distinctive among individuals. A detected pattern of individual distinctiveness in *C. mentalis* would not only add to the literature on the little-studied suboscine birds, but such a pattern may yield new methodological possibilities for using vocalizations to rapidly assess male abundance, helpful for conservation efforts.

## METHODS

### Study species

*C. mentalis* males compete for females by performing elaborate dances with visible and audible displays at specific locations, called “leks”. *C. mentalis* exhibits a condensed exploded lek, where three to 10 males engage in uncoordinated, non-territorial display for females (Skutch 1949). Leks occur during the dry season and at least two months of the rainy season (Kricher and Davis 1998). Furthermore, males exhibit lek fidelity, often remaining with the same lek for multiple years (Kricher and Davis 1998). Displays consist of numerous vocalizations, mechanical sounds, and motions. The whistle-peep call (described as “*Psit psit psit p’tsweeee - - psip*” by Skutch 1949) appears to be a contact call between males and is used frequently when females or green-plumage juvenile males are not present at the lek. We used this call to test for individual vocal distinctiveness among males across four sites, spanning a stretch of 57 km across the Isthmus of Panama (Fig 1).

**Figure 1:**
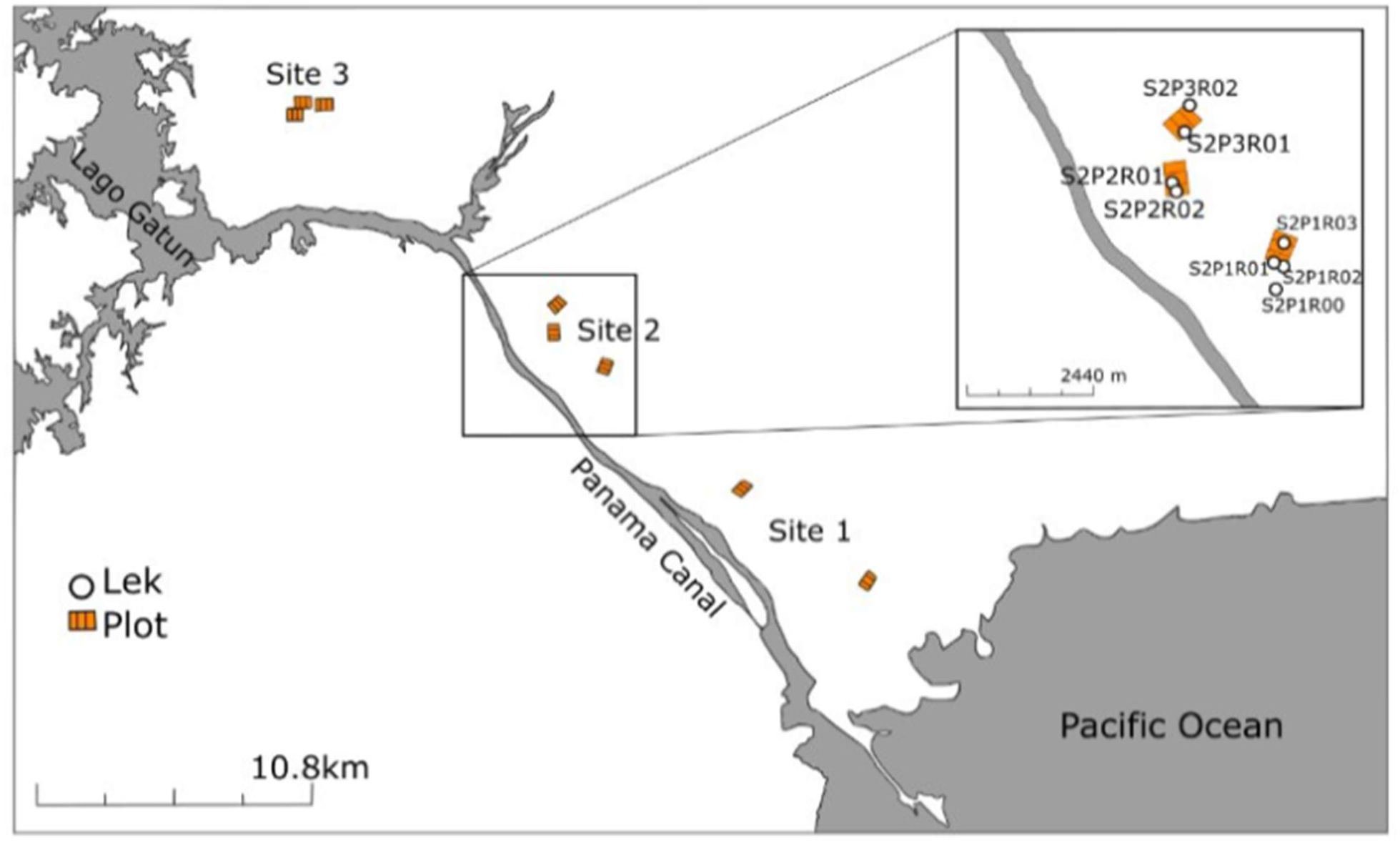
Map of the lower half of the Isthmus of Panama with sites marked. Each orange box symbolizes a plot, with white dots denoting a lek. Inlay is a closer image of the plots at Site 2 with labeled leks. In the lek labels, “S” refers to site, “P” to plot, and “R” to lek. Adapted from Google Earth imagery and GPS points taken by the authors.

### Acoustic recordings

Behavioral observation sessions at each lek were recorded during the breeding seasons in March-June 2017, May-June 2018, and January 2019. During each observation, a focal male was selected, and an observer recorded both verbal annotation of behaviors and the sounds of the lek into a handheld Tascam DR-05 Linear PCM recorder (at 44.1kHz sampling rate, 16-bit mono) for a minimum of 20 minutes. Focal males were identified if banded. We did not repeat sampling at a lek in any given year to avoid duplicating observations of unbanded males.

### Individual selection

To assess individual distinctiveness in male vocalizations, we extracted a total of 120 whistle-peep calls from recordings of calls made by 17 males at 13 leks. Recordings and males were selected based on recording quality (high signal to noise ratio) as well as the number of whistle-peep calls made by each male during the 20-minute recording period. For our dataset, we only retained males with at least three recorded whistle-peeps (mean = 6.9 whistle-peeps per male; range = 3-30).

### Acoustic traits

From each whistle-peep, we extracted nine acoustic variables related to peep quantity, temporal measurements of call and silence intervals, and frequency measurements of all call components (Fig 2). The timing of each call component (peep, whistle, and explosive peep) and the silence between each sound, as well as the low, high, maximum/peak, center, and delta frequencies were quantified using RAVEN Pro version 1.5 (Bioacoustics Research Program 2011) and extracted from the same program by a single author (JTT) (see Fig 2 for call components and the definition of each measurement). We also derived mean, minimum, and maximum values from each acoustic trait and included these metrics in the analysis.

**Figure 2.**
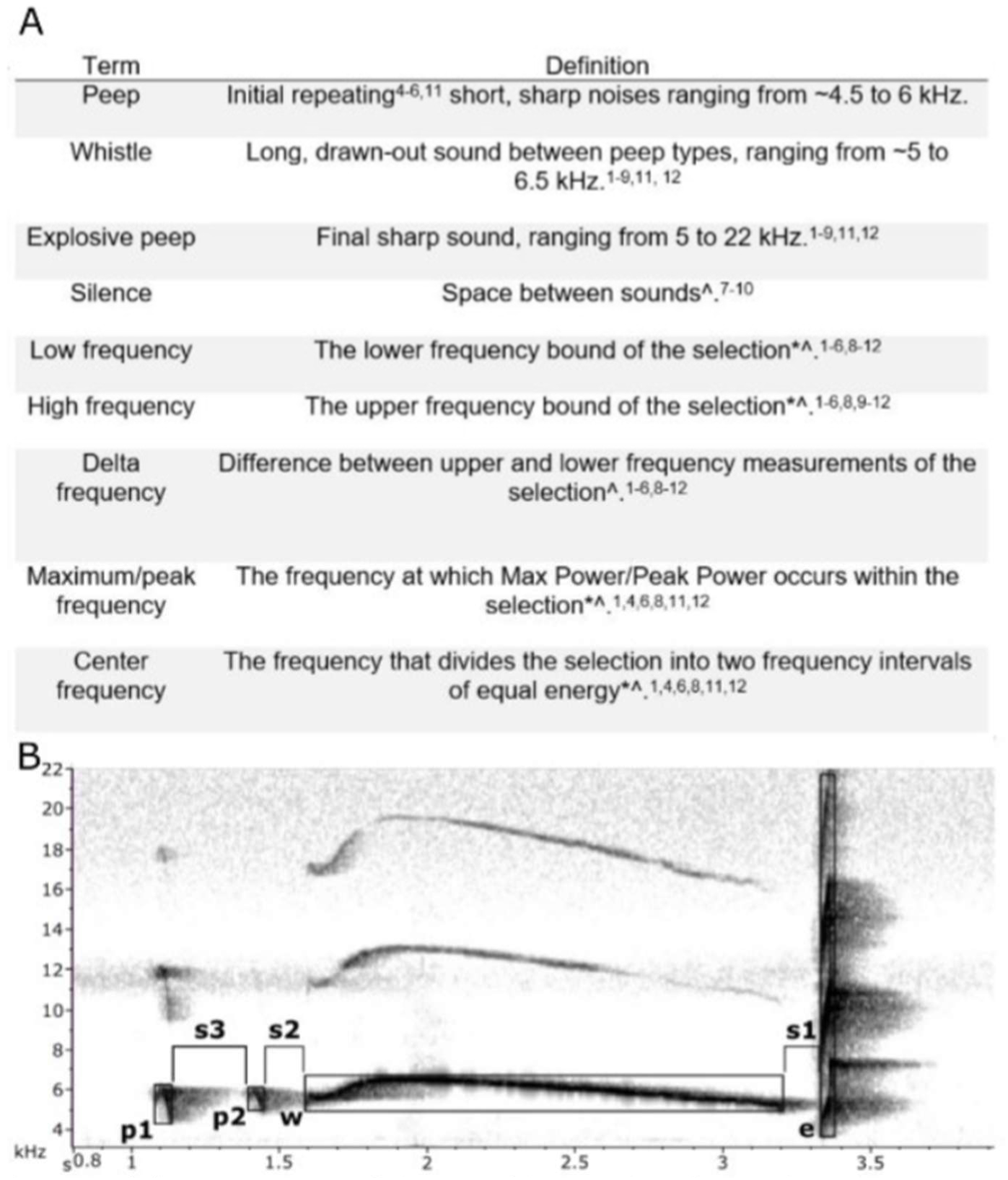
(A) Descriptions of the variables used in the analyses. Definitions marked with * are directly quoted from the manual of RAVEN Pro (Bioacoustics Research Program 2011). Definitions marked with ^A^ refer to measurements that are dependent on the selection window. Superscripts indicate references to the use of each variable in similar studies as follows: ^1^ Bard et al. 2002;^2^ Cooper and Cuervo 2017;^3^ Fitzsimmons et al. 2008; ^4^ Foote et al. 2013;^5^ García et al. 2018; ^6^lppi et al. 2011; ^7^ Isler et al. 2005; ^8^Kirschel et al. 2011;^9^ Lovell and Lein 2004; ^10^ Lovell and Lein 2013; ^11^ Moseley and Wiley 2013; ^12^ Sementili-Cardoso et al. 2018; ^13^Van Doren et al. 2017; ^14^ Wilson and Mennill 2010. Superscripts for the whistle and explosive peep definitions refer to the use of the duration variable in similar studies. (B) Sample whistle-peep sonogram produced by RAVEN Pro (Bioacoustics Research Program 2011). p refers to peep, w refers to whistle, e refers to explosive peep, and s refers to silence. For each sonogram, peeps were labeled pl, p2, etc. in accordance with proximity to the whistle syllable; silence is labeled similarly in accordance with proximity to the explosive peep syllable.

### Statistical analyses

To assess whether the measured acoustic traits of whistle-peep calls could be used to accurately identify individual birds, we conducted a flexible discriminant analysis (hereafter FDA). FDAs are commonly used when analyzing individual variation in birds, and the analysis is favored for its ability to analyze a combination of song variables. Furthermore, it is thought to mirror the algorithm used by birds when discriminating between familiar and unfamiliar conspecifics (Clapperton and Jenkins 1987, Bard et al. 2002, Kirschel et al. 2012). To identify discriminant functions that optimally separate groups within the FDA, we used polynomial regression as the optimal scaling algorithm. We used a jackknife, or leave-one-out, approach (Manly 1994), in which one song was retained for validation via assessment of classification accuracy, while all other songs were used for model training. This model yielded a binary classification for each song (1=correctly classified to individual, 0=classified to another individual). We implemented the FDA in Program R (R Core Team 2019) using the *mda* package (Hastie and Tibshirani 2017) and custom randomization and bootstrapping scripts.

Because our dataset was unbalanced (different number of calls per male), we estimated overall classification accuracy by modeling the binary classification using a binomial generalized linear mixed model (GLMM) (intercept only) with individual identity as a random effect. We used the FDA model to calculate discriminant variables, from which we calculated variable loadings as the correlation between these discriminant variables and each original song trait (Supplementary Information). Song traits that cumulatively contribute to at least 90% of the total variance of each discriminant function are considered maximally important. To assess if the FDA-derived discriminability is greater than random (if calls were randomly distributed between individuals), we ran 500 bootstrap replicates of each FDA using a randomized dataset. We then compared the overall classification accuracy, a single point value, to these bootstrapped distributions.

## RESULTS

### Flexible discriminant analysis

In total, nine dimensions in the FDA had an eigenvalue > 1, accounting for 98.2% of the total variance (Table 1). The first three discriminant dimensions explained >90% of the variance in the acoustic trait dataset, with the first dimension explaining 77.83%, the second dimension explaining 7.66%, and the third dimension explaining 4.59% of total variance (Table 1). Six additional dimensions, each accounting for <3% of the variance, accounted for an additional 8.14% of total variance. The first dimension was correlated most highly (r>|0.3|) to the number of peeps, and both time and frequency traits, including 18 acoustic traits (seven temporal and 10 frequency traits). The second dimension included only four acoustic traits (one temporal trait and three frequency traits) with r>|0.3|, whereas the third dimension correlated most strongly (r > |0.3|) with 14 acoustic traits (all frequency traits).

**Table 1.** Summary of loadings from the individual FDAs. Each dimension refers to a variable describing some amount of the variance as expressed in the explained variance column. Cumulative explained variance (%) is shown in the last column. Rows in bold refer to dimensions that contribute to >96% of total variance (eigenvalue > 1).

| Dimension | Eigenvalue | Explained<br>variance (%) | Cumulative<br>explained<br>variance (%) |
| --- | --- | --- | --- |
| **1** | **121.84** | **77.83** | **77.83** |
| **2** | **12** | **7.66** | **85.49** |
| **3** | **7.18** | **4.59** | **90.08** |
| **4** | **3.98** | **2.54** | **92.62** |
| **5** | **2.57** | **1.64** | **94.26** |
| **6** | **2.37** | **1.51** | **95.77** |
| **7** | **1.42** | **0.91** | **96.68** |
| **8** | **1.23** | **0.79** | **97.47** |
| **9** | **1.16** | **0.74** | **98.21** |
| 10 | 0.8 | 0.51 | 98.72 |
| 11 | 0.67 | 0.43 | 99.15 |
| 12 | 0.41 | 0.26 | 99.42 |
| 13 | 0.39 | 0.25 | 99.67 |
| 14 | 0.28 | 0.18 | 99.85 |
| 15 | 0.16 | 0.1 | 99.95 |
| 16 | 0.08 | 0.05 | 100 |

### Individual distinctiveness

At the population level and without accounting for the uneven sampling among individuals, the FDA correctly classified 85.7% of the calls to the correct individual male. After accounting for the unbalanced dataset using a GLMM, the estimated population-level classification accuracy was 81.1% (95% CI: 63.1-91.5%; range: 16.7-100%). We found our observed population-level classification accuracy to fall outside of the confidence interval of the expected population-level classification accuracy (random classification accuracy based on the randomized FDA = 5.47%, 95% CI: 4.47-6.67%; Table 2). Of the 17 males in this study, 12 males were identified via vocal traits with >75% accuracy and all males had higher classification accuracy than random (Fig 3).

**Table 2.**
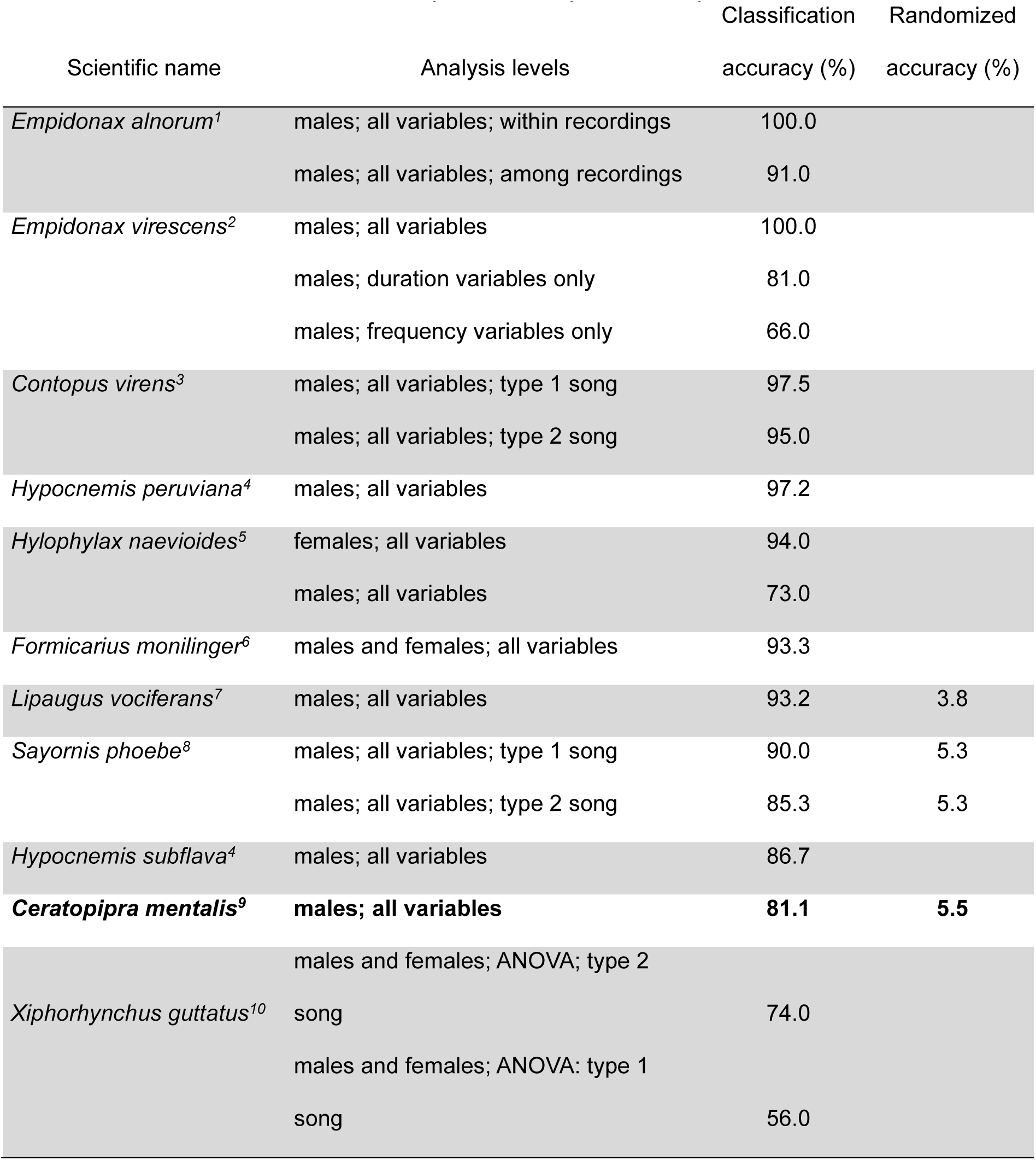
Compilation of existing suboscine studies that also used a discriminant function analysis (unless otherwise noted) and their findings, organized in descending order by species from the highest to lowest classification. Randomized accuracy is given when included in a study. Results from this study are presented in bold. Superscripts indicate references as follows: ^1^Lovell and Lein 2004; ^2^Wiley 2005; ^3^Clark and Leung 2011; ^4^Seddon and Tobias 2010; ^5^Bard et al. 2002; ^6^Kirschel et al. 2011; ^7^Fitzsimmons et al. 2008; ^8^Foote et al. 2013; ^9^This study; ^10^Moseley and Wiley 2013.

**Figure 3.**
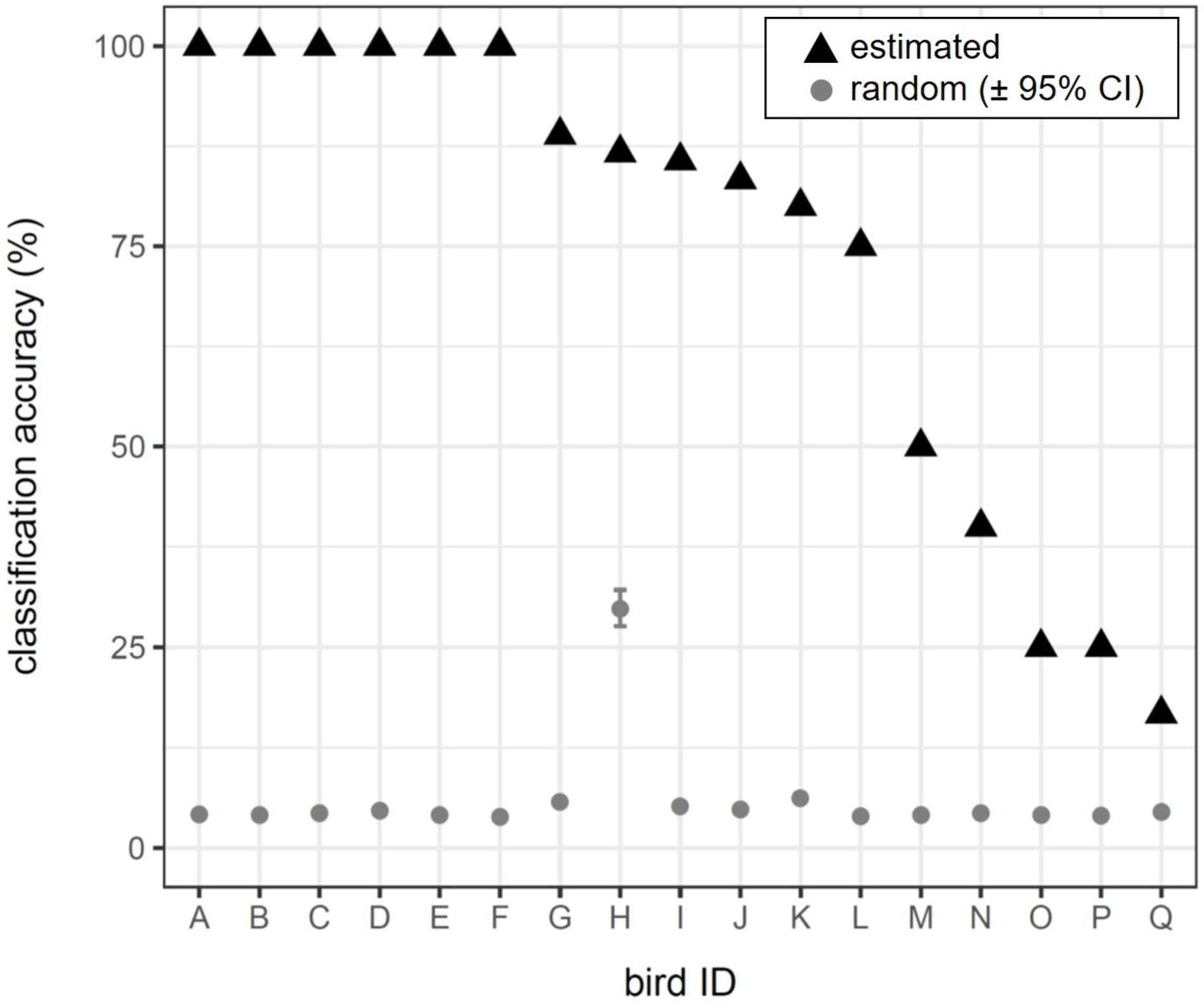
Graph of observed classification accuracy for each individual male, contrasted with random classification values.

We compared our population-level results to other studies of suboscine birds, including only those that used FDA to classify vocalizations. We found that our classification accuracy is comparable to other studies of suboscines (Table 2).

## DISCUSSION

Our examination of the whistle-peep calls of male *C. mentalis* provides the first known evidence of individual distinctiveness in manakins (Family Pipridae). Our estimate of distinctiveness, defined as correct classification to an individual, fits within the range of estimates presented in other studies of suboscine birds (Table 2). These findings add to a growing body of literature indicating that the general absence of song learning and less complex syrinxes of suboscines does not preclude individual distinctiveness in their vocalizations (Bard et al. 2002, Kroodsma 2013, Payevsky 2014). Our findings of individual distinctiveness open up the possibility that individuals may be able to use vocalizations to discriminate among conspecifics, and for use in mate choice and male-male competition. Nevertheless, further research is needed to assess the functions of individual distinctiveness in manakins and other suboscines.

Research on variation in suboscine vocalizations has focused largely on territorial species to date, including *Hylophylax naevioides*, *Empidonax alnorum*, *Contopus virens*, *Formicarius moniliger*, and *Sayornis phoebe*, to name a few (Bard et al. 2002, Lovell and Lein 2004, Clark and Leung 2011, Kirschel et al. 2012, Foote et al. 2013). Here, we found similar evidence for individual vocal distinctiveness in a non-territorial lekking species, suggesting that delineated territories are not a prerequisite for individual vocal distinctiveness.

In lekking species, individually distinct vocalizations may be particularly important in female attraction or to convey status given the high mating skew and established dominance hierarchy observed in many of these species (Lambrechts and Dhondt 1995, Naguib and Riebel 2006, DuVal et al. 2018). Vocalizations of various bird species are known to convey information regarding hierarchical status, individual origin, age, gender, motivation, and quality (Clapperton and Jenkins 1987, Lambrechts and Dhondt 1995, Naguib and Riebel 2006). Evidence for vocally transmitted information use by females for mate selection exists in at least one species of manakin (Long-tailed Manakin [*Chiroxiphia linearis*], Trainer and McDonald 1995).

Nevertheless, studies of sexual selection and mate choice in female manakins rarely assess the role of vocalizations. Furthermore, the role that vocalizations may play in maintaining the dominance hierarchy within exploded leks of male manakins is unclear. The individual distinctiveness of male *C. mentalis* suggests that males are communicating information about themselves via these commonly produced contact calls. Further study of what information is being transmitted will be fruitful for studies of sexual selection and behavior.

Manakins are essential to the health and persistence of Neotropical forests as important seed dispersers (Poulin et al. 1999). They are sensitive to climate change and are expected to decline over the next 50 years, with 20% of manakins predicted to become extinct in their current range and half of all species expected to lose 80% of their current range (Anciães and Peterson 2006, Brawn et al. 2017). Such declines threaten the well-being of lowland Central American forests. *C. mentalis* is predicted to experience a 49% range contraction over the next several decades (Anciães and Peterson 2006). The presence of individual distinctiveness in *C. mentalis* vocalizations allows for the possibility of establishing a system of remote acoustic monitoring of individuals based on these contact calls, leading to robust, non-invasive population methods to assess changes in male abundance. Remote acoustic monitoring may serve as an alternative to mark-recapture and point-count studies in places that are hard to survey or where rapid assessments are needed (Katzner et al. 2011). However, further work is needed to compare estimates of male abundance based on these different techniques and to examine vocal consistency across breeding seasons to establish a robust method for non-invasive acoustic population monitoring.

In conclusion, we found individual distinctiveness in the whistle-peep calls of *C. mentalis* males, a lekking suboscine. This work adds to the growing body of literature challenging the assumption that suboscines lack individual vocal distinctiveness. We argue for the need to conduct more intensive study on the functions of suboscine vocalizations and the potential to use vocalizations for non-invasive population monitoring of *C. mentalis*, and potentially other suboscines.

## ACKNOWLEDGEMENTS

We would like to acknowledge the University of Wyoming and the Wyoming Research Scholars Program for funding this project, as well as the Smithsonian Tropical Research Institute (STRI) for logistical support. We thank Dan Albrecht-Mallinger, Alex Bartolo, and Libby Megna for their assistance in the field. All research was conducted under the auspices of permits provided by the Ministerio del Ambiente and approved under the Institutional Animal Care and Use committees at STRI and the University of Wyoming.

