## Supplemental Tables for "Males of a suboscine bird (*Ceratopipra mentalis*) exhibit individual distinctiveness in their contact call"

**Supplemental Table S1.** Description of the acoustic variables measured and included in the analysis. Variables with "Duration" in the name refer to timing traits, and variables with "Frequency" refer to frequency traits. As each whistle-peep contains a variable number of peeps, we included only the minimum, maximum, and mean values of all duration and frequency variables for all peeps in a given call. "Abbrev" is the abbreviation of the acoustic variable and call component, as it appears in the data set and loadings.

| Acoustic Variable | Call Component | Abbrev | Min | Mean | Max | Variance |
|---|---|---|---|---|---|---|
| Number of peeps | | NumPeeps | 2 | 3.49 | 7 | 0.0869 |
| Sound duration | peep (minimum) | Tp_min | 0.03 | 0.11 | 0.17 | 0.0662 |
| | peep (maximum) | Tp_max | 0.05 | 0.11 | 0.23 | 0.0661 |
| | peep (mean) | Tp_mean | 0.04 | 0.11 | 0.19 | 0.0571 |
| | whistle | T_w | 1.07 | 1.7 | 2.6 | 0.0277 |
| | explosive peep | T_e | 0.04 | 0.17 | 0.26 | 0.2865 |
| Silence duration | inter-peep (minimum) | Ts_min | 0.02 | 0.11 | 0.23 | 0.1597 |
| | inter-peep (maximum) | Ts_max | 0.1 | 0.22 | 0.67 | 0.1136 |
| | inter-peep (mean) | Ts_mean | 0.07 | 0.16 | 0.43 | 0.086 |
| | pre-explosive peep silence | T_s1 | 0.08 | 0.39 | 0.69 | 0.1234 |
| Low frequency | peep (minimum) | L_min | 3322.6 | 4094.12 | 4800 | 0.0063 |
| | peep (maximum) | L_max | 3564.2 | 4454.84 | 5134.9 | 0.0052 |
| | peep (mean) | L_mean | 2609.76 | 4267.4 | 4865.77 | 0.0068 |
| | whistle | L_w | 3624.7 | 4100.29 | 5134.9 | 0.0049 |
| | explosive peep | L_e | 1208.2 | 3489.45 | 4668.5 | 0.0353 |
| High frequency | peep (minimum) | H_min | 4470.4 | 6167.76 | 6524.4 | 0.0052 |
| | peep (maximum) | H_max | 5014.1 | 6344.51 | 6826.4 | 0.0038 |
| | peep (mean) | H_mean | 3842.14 | 6202.73 | 6584.8 | 0.0055 |
| | whistle | H_w | 6222.3 | 6895.34 | 7551.4 | 0.0012 |

| | | | | | | |
|---|---|---|---|---|---|---|
| | explosive peep | H_e | 11506.8 | 21770.86 | 21989.6 | 0.0219 |
| Maximum frequency | peep (minimum) | M_min | 4306.6 | 5646.37 | 6029.3 | 0.0057 |
| | peep (maximum) | M_max | 4651.2 | 5986.89 | 6373.8 | 0.0038 |
| | peep (mean) | M_mean | 3565.9 | 5781.14 | 6115.45 | 0.0058 |
| | whistle | M_w | 5426.4 | 6301.45 | 6976.8 | 0.0022 |
| | explosive peep | M_e | 4995.7 | 9857.87 | 11369.5 | 0.0446 |
| Center frequency | peep (minimum) | C_min | 4220.5 | 5620.66 | 5943.2 | 0.0045 |
| | peep (maximum) | C_max | 4737.3 | 5927.35 | 6287.7 | 0.0035 |
| | peep (mean) | C_mean | 3514.22 | 5739.5 | 6029.3 | 0.0054 |
| | whistle | C_w | 5598.6 | 6192.06 | 6804.5 | 0.0013 |
| | explosive peep | C_e | 7752 | 10170.82 | 10680.5 | 0.0024 |
| Difference between high and low frequency | peep (minimum) | D_min | 543.7 | 1746.71 | 2476.8 | 0.0935 |
| | peep (maximum) | D_max | 906.2 | 2167.86 | 2960.1 | 0.057 |
| | peep (mean) | D_mean | 845.75 | 1968.32 | 2567.45 | 0.0594 |
| | whistle | D_w | 1510.3 | 2815.6 | 3262.2 | 0.0241 |
| | explosive peep | D_e | 6838.4 | 18254.56 | 20660.5 | 0.0468 |

**Supplemental Table S2.** Loadings of the FDFA performed for individual variation. Loadings >|0.3| are shown in bold. For clarity, only the first 12 dimensions are shown, even though the first three explain 90% of the variance in the vocal trait dataset.

| | Discriminant dimension | | | | | | | | | | | |
|---|---|---|---|---|---|---|---|---|---|---|---|---|
| **Variable** | **1** | **2** | **3** | **4** | **5** | **6** | **7** | **8** | **9** | **10** | **11** | **12** |
| NumPeeps | **-0.46** | 0.00 | 0.12 | 0.12 | 0.10 | -0.05 | -0.29 | -0.17 | 0.08 | -0.07 | -0.03 | -0.25 |
| Tp_min | **0.74** | -0.18 | 0.00 | -0.15 | -0.12 | 0.08 | 0.07 | 0.14 | -0.01 | **0.37** | -0.05 | -0.19 |
| Tp_max | **0.53** | -0.25 | -0.13 | -0.10 | -0.12 | 0.01 | -0.21 | 0.06 | 0.08 | **0.33** | -0.07 | -0.07 |

| | | | | | | | | | | | | |
|---|---|---|---|---|---|---|---|---|---|---|---|---|
| Tp_mean | **0.72** | -0.23 | -0.09 | -0.13 | -0.11 | 0.02 | -0.07 | 0.10 | 0.05 | **0.39** | -0.08 | -0.13 |
| Ts_min | **-0.49** | 0.20 | 0.16 | -0.11 | **-0.34** | 0.27 | 0.15 | -0.10 | 0.02 | -0.25 | 0.01 | 0.13 |
| Ts_max | -0.21 | 0.07 | 0.06 | -0.05 | -0.27 | -0.02 | -0.10 | -0.13 | -0.25 | -0.15 | -0.28 | -0.13 |
| Ts_mean | **-0.37** | 0.15 | 0.11 | -0.13 | **-0.40** | 0.07 | -0.03 | -0.17 | -0.19 | -0.26 | -0.24 | -0.06 |
| T_w | 0.20 | **0.80** | 0.10 | **-0.38** | -0.20 | -0.16 | 0.10 | -0.16 | -0.02 | -0.06 | -0.10 | 0.06 |
| T_s1 | **0.48** | 0.10 | 0.08 | -0.13 | **-0.33** | -0.20 | -0.09 | 0.07 | **-0.45** | 0.06 | 0.27 | 0.25 |
| T_e | **0.93** | -0.17 | -0.13 | 0.01 | -0.08 | -0.05 | 0.04 | -0.07 | 0.05 | -0.15 | 0.05 | -0.02 |
| L_min | -0.02 | 0.04 | **-0.67** | 0.04 | 0.01 | -0.25 | 0.21 | 0.06 | 0.01 | -0.02 | 0.10 | 0.14 |
| L_max | 0.15 | -0.12 | **-0.68** | -0.08 | -0.06 | -0.29 | 0.05 | 0.08 | 0.18 | 0.06 | 0.07 | 0.15 |
| L_mean | 0.08 | 0.01 | **-0.61** | 0.05 | 0.01 | **-0.36** | **0.41** | -0.06 | -0.02 | 0.06 | 0.02 | 0.23 |
| L_w | **-0.62** | **0.34** | **-0.51** | 0.18 | 0.06 | -0.08 | -0.17 | 0.12 | 0.07 | -0.05 | -0.11 | -0.04 |
| L_e | **-0.77** | 0.02 | 0.01 | -0.11 | 0.21 | -0.06 | 0.15 | **0.37** | 0.00 | -0.01 | 0.16 | -0.09 |
| H_min | **0.45** | 0.19 | **-0.51** | **-0.37** | 0.11 | **0.36** | -0.02 | 0.04 | -0.16 | -0.03 | 0.05 | 0.08 |
| H_max | 0.27 | 0.14 | **-0.59** | -0.20 | 0.30 | **0.38** | -0.14 | -0.08 | -0.10 | -0.07 | 0.20 | 0.03 |
| H_mean | **0.37** | 0.18 | **-0.44** | -0.23 | 0.27 | 0.26 | 0.19 | -0.18 | -0.22 | 0.00 | 0.04 | 0.15 |
| H_w | 0.22 | **0.41** | -0.28 | **0.44** | -0.30 | **0.32** | -0.05 | 0.14 | -0.09 | 0.03 | 0.17 | 0.04 |
| H_e | **0.97** | -0.02 | -0.04 | 0.15 | 0.10 | 0.07 | 0.01 | -0.02 | 0.00 | 0.01 | -0.04 | 0.00 |
| M_min | 0.29 | 0.21 | **-0.54** | -0.27 | 0.22 | **0.32** | 0.06 | 0.04 | -0.14 | -0.02 | 0.04 | 0.08 |
| M_max | 0.18 | 0.19 | **-0.62** | -0.17 | **0.35** | 0.30 | 0.00 | -0.11 | -0.16 | -0.10 | 0.16 | -0.07 |
| M_mean | 0.27 | 0.19 | **-0.44** | -0.16 | **0.32** | 0.21 | 0.24 | -0.19 | -0.23 | 0.00 | 0.03 | 0.12 |
| M_w | -0.10 | 0.19 | -0.27 | **0.49** | -0.26 | 0.25 | -0.06 | 0.04 | -0.19 | 0.08 | 0.10 | -0.02 |
| M_e | -0.21 | 0.16 | -0.19 | 0.06 | 0.12 | -0.03 | 0.11 | 0.04 | 0.25 | -0.02 | **0.38** | 0.03 |
| C_min | **0.33** | 0.23 | **-0.57** | **-0.32** | 0.20 | **0.31** | 0.04 | -0.03 | -0.13 | 0.01 | 0.05 | 0.10 |
| C_max | 0.21 | 0.18 | **-0.64** | -0.18 | 0.30 | **0.32** | 0.02 | -0.17 | -0.14 | -0.08 | 0.19 | -0.03 |
| C_mean | 0.27 | 0.19 | **-0.46** | -0.18 | 0.30 | 0.20 | 0.27 | -0.25 | -0.22 | 0.01 | 0.04 | 0.15 |
| C_w | -0.10 | 0.22 | **-0.51** | **0.60** | -0.28 | 0.19 | -0.08 | -0.07 | -0.06 | 0.15 | 0.22 | -0.07 |
| C_e | -0.16 | **0.32** | -0.21 | 0.16 | -0.03 | 0.17 | 0.20 | 0.30 | 0.09 | -0.04 | **0.35** | 0.07 |
| D_min | **0.38** | 0.23 | -0.08 | -0.26 | 0.10 | **0.52** | -0.07 | -0.03 | **-0.33** | -0.19 | -0.04 | -0.02 |

| | | | | | | | | | | | | |
|---|---|---|---|---|---|---|---|---|---|---|---|---|
| D_max | 0.29 | 0.12 | 0.04 | -0.24 | 0.29 | **0.60** | **-0.35** | -0.12 | -0.07 | 0.01 | 0.10 | -0.07 |
| D_mean | **0.34** | 0.19 | 0.03 | -0.25 | 0.25 | **0.60** | -0.16 | -0.15 | -0.22 | -0.07 | 0.02 | -0.04 |
| D_w | **0.70** | -0.03 | 0.27 | 0.14 | -0.26 | 0.29 | 0.12 | -0.01 | -0.13 | 0.07 | 0.22 | 0.06 |
| D_e | **0.97** | -0.02 | -0.04 | 0.15 | 0.05 | 0.07 | -0.02 | -0.08 | 0.00 | 0.01 | -0.07 | 0.01 |
| eigenvalue | 121.84 | 12.00 | 7.18 | 3.98 | 2.57 | 2.37 | 1.42 | 1.23 | 1.16 | 0.80 | 0.67 | 0.41 |
| explained var. (%) | 77.83 | 7.66 | 4.59 | 2.54 | 1.64 | 1.51 | 0.91 | 0.79 | 0.74 | 0.51 | 0.43 | 0.26 |
| cumulative var. (%) | 77.83 | 85.49 | 90.08 | 92.62 | 94.26 | 95.77 | 96.68 | 97.47 | 98.21 | 98.72 | 99.15 | 99.42 |
